# Late Quaternary climate-driven shifts in arctic plant distributions

**DOI:** 10.64898/2025.12.10.693528

**Authors:** Paul Markley, Barnabas H Daru

**Affiliations:** Stanford University

**Keywords:** Arctic, climate, distributions, glaciation, quaternary, refugia

## Abstract

**Aim:** Glaciation events shaped the present distribution of many plants and their biodiversity in the northern hemisphere. Glacial expansion forced many species south and came with much colder global temperatures, while glacial recession brought warmer temperatures and newly colonizable land without competition. However, the changes in plant diversity associated with glacial retreat and the ensuing climatic changes is not well understood. In this study, we quantify Late Quaternary climate-driven changes in Arctic plant diversity by integrating climatic shifts in species distributions since the last glacial maximum (20-16 kya) and mid-Holocene (5 kya) across the circumpolar arctic.

**Location:** The geographic arctic, 66°N.

**Taxon:** Vascular plants.

**Methods:** We built species distribution models using phyloregion v. 1.0.9 in R using occurrence data from the Global Biodiversity Information Facility and climate rasters from Worldclim v2.1 for the present and v1.4 for the mid-Holocene and Last Glacial Maximum.

**Results and Discussion:** We found limited evidence for decreases in weighted endemism, and species richness along with diverging north-south shifts in the centroids of many distributions contrary to expectations of increased alpha diversity since the last glacial maximum. Decreases in species alpha diversity, while already quite low in the arctic, may be reflective of an increasingly variable arctic climate that disfavors plants with a slow dispersal ability. This is especially important given the projected increase in global temperature across many shared socioeconomic pathway scenarios and can be contrasted with our results of the Mid-Holocene, which was roughly a degree warmer than it is today. The arctic is presently warming at roughly two to five times the rate of the mid-latitudes and equator and understanding how plants have responded in the past will help inform on how they may change in the future.

## Introduction

Climatic changes since the Last Glacial Maximum (LGM, 20-16kya) [1. Birks 2008; 2. Alsos et al. 2009] have shaped the present distribution of arctic plants in the Northern Hemisphere [3. Markley et al. 2024]. Global temperatures at the LGM are estimated to be 5°C cooler than present-day [4. Shakun and Carlson 2010; 5. Osman et al. 2021], leading to tundra plant distributions that extended farther south than their current ranges [6. Hanberry 2024; 7. Hanberry 2023]. For many species at high latitudes, much of the land today considered to be arctic would have been covered by ice sheets that likely would have denied a continuous presence in these areas leading to plant population centers at lower latitudes [8. Bigelow et al. 2003; 9. Pellissier et al. 2016]. These distributions are expected to have been lower in elevation than present-day because of growing ice cap glaciers at mountain peaks that prevented the persistence of most high elevation or alpine plant populations [10. Binney et al. 2017], which usually are either the same species found in the arctic or are at least close relatives or hybrids within the same genus [11. Ebersbach et al. 2020; 12. Boulanger-Lapointe et al. 2014]. Glaciation fragmented the southward shifting plant populations into refugial sites like Beringia, the land between Siberia and Alaska [13. Hultén 1968], and other smaller nunatak refugia, which are small rocky outcroppings at high elevations that may have remained unglaciated in mountain ranges [14. Abbott and Brochmann 2003; 15. Hoffecker et al. 2014]. These refugial sites for arctic species can be considered akin to sources or hotspots of diversity for plants of high latitudes and elevations, and with time these hotspots may have moved or changed in size as climate shifted individual species ranges [1–4, 11, 14].

Previous studies have proposed that arctic plant distributions generally shifted north at the end of the LGM as global temperatures rose, colonizing land that had been otherwise occupied entirely by glaciers from well documented refugia like Beringia, but possibly from cryptic refugia like nunataks [16. DeChaine 2008; 17. Oke et al. 2023; 18. Anderson and Lozhkin 2015]. The strongest direct evidence of historical plant presence in any given area of the arctic is typically through fossil data such as charcoal or pollen [8. Bigelow et al. 2003; 18. Anderson and Lozhkin 2015] while distributional shifts and colonization events may be inferred through statistical models or genetic data [9. Pesselier et al. 2016; 19. Strong 2023; 20. Eidesen et al. 2013]. Previous studies on plant range expansions into and within the arctic propose that arctic plants dispersed post-LGM across the Atlantic Ocean and the Bering Strait [14. Abbott and Brochmann 2003; 20. Eidesen et al. 2013; 21. Gussarova et al. 2015]. The frequency of these events led to circumpolar and montane disjunct distributions [3. Markley et al. 2024; 13. Hultén 1968]. Many plant communities of the arctic are relatively species poor when compared to lower latitudes, most likely because of the prediction of decreasing species richness and increasing range sizes with latitude observed globally today [22. Liu et al. 2025; 23. Brodie and Mannion 2023; 24. Guo et al. 2022; 28. Daru 2024]. While changes in arctic plant distributions with climate from the LGM to the present have been confirmed primarily through fossil data, these sources may not necessarily capture the full extent of these species ranges at the LGM as predicted through species distribution modeling (SDM) since they frequently come from only one soil core, usually from lake sediments, limiting inference of the possible extent of the plant’s true occurrence at a given time [25. Gavin et al. 2014; 26. Svenning et al. 2008].

Projecting backwards in time or hindcasting arctic plant distributions can show how plants have responded to shifts in climate coming out of the LGM [1. Birks 2008; 25. Gavin et al. 2014]. SDMs can enable research on arctic plant distributions under any climate scenario by using known presences and predicting into unsurveyed regions [27. Panchen et al. 2019]. SDMs can therefore provide high resolution range maps for many species across the globe which complements genetic and paleobiological studies on the past distributions of organisms [25, 26]. Modeling past species distributions using present data and validating them using fossil data may reveal historical patterns of plant diversity and give additional context to projected and ongoing anthropogenic climate change. We hypothesize that the climatic transition from the LGM to the Mid-Holocene (5 kya) and the present is comparable to the projected climatic trends [2, 25–26]. Thus, aggregating SDMs to create species richness maps can reveal hidden diversity centers [28. Daru 2024], which although the arctic may be low in unique species they are still likely to detect hidden plant diversity since many plants share a common geographic and evolutionary origin given the geological history of glaciation in the area [1,3,13–14,20]. The hindcast SDMs can also enable analysis of how ranges could have shifted out of the LGM and into the present, which can be relevant for understanding patterns of species migrations in a warming climate [25–26, 29. Naughtin et al. 2024].

Anthropogenic climatic change, particularly higher global temperatures, is projected to shift many plant distributions poleward and higher in elevation [30. Niskanen et al. 2019; 31. Boisvert-Marsch et al. 2014]. A well-documented phenomenon is the northward shifting boreal-tundra ecotone whereby the plants of lower latitudes colonize continually farther north as temperature no longer limits recruitment and survival of the plants that compose the boreal forest community [31]. Current climatic changes within the arctic are proceeding at nearly five times the rate of anywhere else on Earth [32. Previdi et al. 2021; 33. Koenigk et al. 2020; 34. England et al. 2021]. The implication is that available land for cold-adapted arctic plants is rapidly dwindling as plant populations along the southern edges of their ranges decline because of increased temperature stress and competition from lower latitudes [35. Lesica and Crone 2017]. Studies on projected climate change using species distribution models have documented the shrinkage of habitable land for plants on the species range edge [35; 36. Kaplan and New 2006]. For many species in the temperate mid-latitudes, this climatic escape into the arctic is limited only by species dispersal ability, or through human assistance via unintentional introductions [37. Wasowicz et al. 2020].

Here, we report changes in arctic plant distributions along latitudes and elevation since the LGM, along with geographical changes in species richness, and the velocity of such changes. We ask: (1) to what extent have arctic plant communities change with respect to range contractions and expansions along latitudes and elevations? (2) which regions experienced the largest changes in species richness? and (3) at what velocities are plant species shifting their ranges in response to climate? We expect that plant ranges shifted north from the LGM to the present, and that certain areas known to have served as refugial sites generally had a higher richness than others. We expect that the velocity of range shifts increased between the LGM and the Mid-Holocene, with the Mid-Holocene to the present showing a faster northward movement. Our findings reveal minimal changes in most Arctic plant species range sizes and elevations, but a consistent upward and outward shift in ranges in those that do. We also demonstrate strong decreases in alpha diversity in the near past, while showing increases since the LGM.

## Materials and Methods

### Species Distribution Models

Plant occurrence data for all plants of the world were downloaded from the Global Biodiversity Information Facility (GBIF) on January 12, 2023 [1. GBIF, 2023; https://doi.org/10.15468/dl.t7mpsk]. This dataset was restricted to plant species found in the Arctic following the species list from Qian et al. (2022) [2]. Calibration areas for species distribution models were created from the species occurrence records using the rangeBuilder R package [3] in R v4.2.2 [4]. We generated alpha hulls around the points and then buffered each polygon based on the estimated phylogenetic dispersal ability of each genus using the castor package functions *fit_sbm_const* and *expected_sbm_distance* [5] with the Smith and Brown megaphylogeny (2018) [6]. Species distribution models were created with the thinned GBIF data, the calibration areas, and climate layers from WorldClim v2.1 [7] at the 5-arcminute resolution for present distributions and WorldClim v1.4 [8] at the 5-arcminute resolution for the LGM and Mid-Holocene using the phyloregion package [9] with the MaxEnt option. All climate rasters from WorldClim were derived from the Model for Interdisciplinary Research on Climate Earth System Model (MIROC-ESM). The climate layers used were mean diurnal range (BIO2), isothermality (BIO3), mean temperature of wettest quarter (BIO8), precipitation of driest month (BIO14), precipitation seasonality (BIO15), precipitation of warmest quarter (BIO18), and precipitation of coldest quarter (BIO19). These were selected based on the variable inflation factor of all 19 bioclimatic variables that were less than five using the usdm package [10]. SDMs were generated using the *sdm* function from the package phyloregion, which uses MaxEnt and five-fold validation to generate suitable distributions of plants. The SDM output included a data table of the model performance computed as Area Under the Curve and True Skill Statistic, and a raster and polygon of the distribution.

### Species Velocities and Centroid Shifts

Species range shift velocities were calculated and analyzed for latitudinal shifts. The magnitude of the velocity was determined through the difference of the distribution’s centroid between time point 1 and time point 2 divided by the number of years spanned between the two for a value in kilometers per year for each species. These values were joined to the polygons and were averaged onto a blank raster of 25 km^2^ and mapped in geographic space. We took the centroids of every distribution and overlaid it onto terrestrial ecorealms of the world and separated each species into two categories which either crossed biogeographical realms such as from the palearctic to the Nearctic or did not across time points. We then assessed the significance of the shifting centroids using the CircStats R package function *v0.test* for within the period shifts alone and *Watson.two* function to compare the shifts between the two time periods [11]. Phylogenetic signals of species range shift velocity were assessed for statistical significance using the phylosignal package [12]. While this package returns statistics frequently used such as K, K*, Cmean, and Pagel’s λ, we considered primarily λ as the primary test statistic with α = 0.05. Values of Pagel’s λ typically range between 0 and 1 where 0 indicates nearly no discernable relationship of the continuous trait and evolution under Brownian motion, and 1 indicates a phylogeny which supports evolution under Brownian motion [19. Pearse et al. 2025].

### Elevational Shifts

To determine whether a significant change in elevation had occurred, we used a spreading dye algorithm. This works by sampling points at random within the geographic distribution and buffering each point iteratively by a set amount until total buffered area is equal to the spatial extent of the input distribution when divided by pi. For every species in this study, we ran the spreading dye algorithm 1000 times and determined the significance by comparing observed elevational shift between two time horizons for a species to the median value of that species across the entire distribution using the digital elevation model from WorldClim v.2 at a 5-minute resolution. P-values were calculated through comparing the number of times the test value was greater or less than the observed value as divided by the total number of runs under a two-tailed test.

### Hotspots of Diversity and Changes in plant alpha diversity

Phylogenetic diversity, phylogenetic endemism, weighted endemism, and richness were calculated using customized functions in the R package phyloregion [9]. Analyses were carried out with grid cells sized at 50 km^2^. Hotspots of diversity metrics were identified as all grid cells that exceeded the 97.5^th^ quantile of all possible values of phylogenetic diversity, weighted endemism, phylogenetic endemism, and richness. We used the *moran.test* and *modified.t.test* functions from the R package SpatialPack [13] to examine the autocorrelation within each raster and to compare between the scenarios for each metric.

### Model Validation: Additional Species Distribution Models

The original raster version of the model estimates for the two past scenarios of the LGM and the Mid-Holocene were converted into a points data frame with the centroid of each pixel as a present occurrence record. This data was then used to generate distribution models of present distributions to verify the accuracy of the hotspot maps and the overlap of the present distributions. We used the same bioclimatic inputs as the original model run for these SDMs. Each species’ modeled predictions (in raster format) were stacked to create a total richness map of Arctic plant species. These richness maps were used to identify biodiversity hotspots which were compared against the two present SDMs and the original present output. In addition to this method of validation, we also considered a scenario where plant species could not disperse in areas covered by glaciers at the LGM in the LGM climate scenario using vector polygon base map from Gowan et al. (2019) [15] to mask raster values that were within the boundaries of that polygon. We ran the SDMs with the original input data described in the first section of the methods.

### Model Validation using machine learning Random Forest

As a final validation of the results for species richness, we used a machine learning algorithm, Random Forest, to examine the aggregated species richness of Arctic flora using the ranger R package (Wright and Ziegler 2015) [16]. We used as input predictors, the bioclimatic raster layers that generated the species distribution models in addition to a raster of the aggregated tetrapod species diversity using range maps from the International Union for the Conservation of Nature (IUCN, 2024) [17] following the methods outlined in Daru (2024). Two spatial kernel density estimates were generated using the spatialEco R package [18]. One was based on the density of input occurrences downloaded from GBIF, while the other assumed equal sampling across the study area. Since there was an additional predictor in the form of mammal richness, we used the *vifstep* function from the usdm package to remove any potentially correlated predictors. We then used the opencv package to create spatial blocks of the world based on known floristic regions and validated the model using five-fold cross validation. Models were tuned by varying the mtry parameter from one to ten, and the model with the lowest root mean square error (mtry = 4) was selected for running random forest.

## Results

### 1. Plant diversity metrics and hotspots

We generated species distribution models for 1,500 plant species of the Arctic under climate scenarios in the present-day versus the LGM (21 kya), and Mid-Holocene (5 kya). Model performance statistics showed a generally higher performance scores across evaluation metrics (Figure S12) indicating the robustness of our predictions. Our modeled range sizes for present distribution of Arctic plant species vary from 24 km^2^ for *Arnica lessingii* Green (Asteraceae) to 27,452,215 km^2^ for *Urtica urens* L. (Urticaceae) with a median range size of 122,870 km^2^ +/-34,557 km^2^.

We explored geographic shifts for common biodiversity metrics including species richness and weighted endemism along with their phylogenetic variants, i.e., phylogenetic diversity and phylogenetic endemism, respectively. We found that areas of highest plant species richness across all three time horizons (present-day, mid-Holocene, and LGM) were in Central Europe and western North America in the Rocky Mountains (Figure 1). A similar pattern was detected for phylogenetic diversity (Figure 1B, F, J). On the other hand, centers of high phylogenetic endemism were detected in Beringia, the Appalachian Mountains, and in northern Caucus Mountains (Figure 1, panels C, G, K). Weighted endemism was consistently low across the Arctic, with highest values in Iceland and Central Europe extending into Scandinavia (Figure 1). These patterns were generally highly clustered spatially based on Moran’s *I* across all metrics and climatic scenarios with an average Moran’s *I* value of 0.8 (p < 0.05 in all cases). Comparing the two past time horizons to the present shows a consistent increase across all metrics in the boreal zone of the Palearctic (Figure S1). There was a decrease in phylogenetic diversity and weighted endemism in higher latitudes of the low and coastal Arctic between both the Mid-Holocene and the present, versus the LGM and the present (Figure S1A, E, C, G).

**Figure 1.**
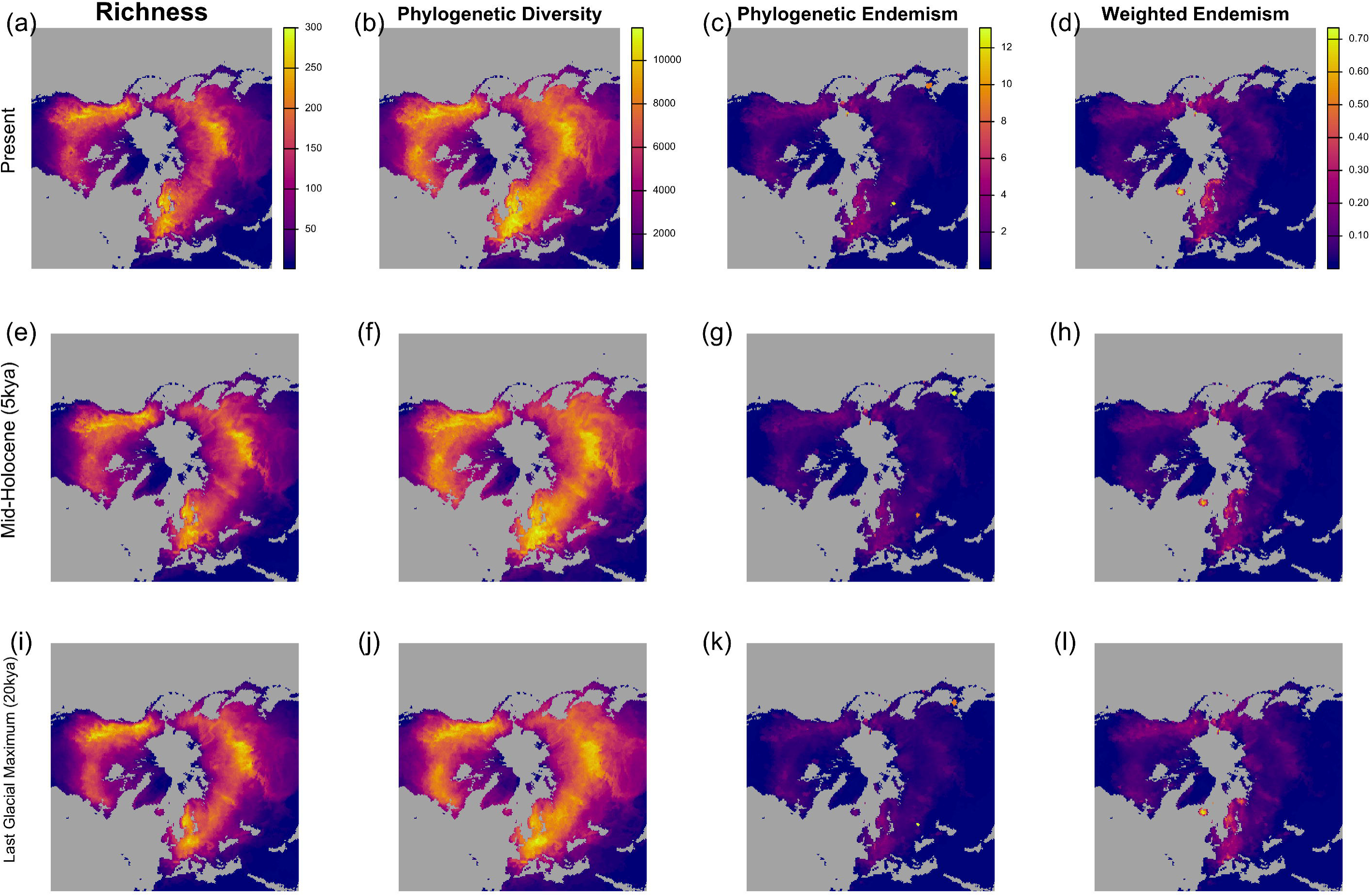
Geographic distributions of Arctic plant diversity. A-D, Present-day diversity metrics, E-H the Mid-Holocene (about 5 kya), and I-L the Last Glacial Maximum (20 kya). The cell values in all cases show higher values in past scenarios. The grids in this raster are roughly 50 km by 50 km projected in EPSG:3996 projection system.

To identify concentrations of these diversity metrics in geographic space, we mapped hotspots of each metric as values in the 2.5^th^ percentile for each metric in each time horizon (Figure 2). Large contiguous hotspots of species richness and phylogenetic diversity in the Arctic are consistent throughout each time horizon to the present. We identified hotspots of these two metrics in the Rocky Mountains of North America, Central Europe, and Mongolia. Phylogenetic endemism and weighted endemism show similar patterns for the Rocky Mountains and Central Europe, but not in Mongolia. Instead, hotspots for these two metrics appear in Iceland and Far East Russia (Figure 2C, 2D). However, we also found losses in hotspots of phylogenetic diversity and species richness toward present-day especially in the Rocky Mountains for species richness and phylogenetic diversity, meaning that these hotspots existed in the past but no longer found today (Figure S1, 2A and 2B). Furthermore, hotspots of weighted endemism and phylogenetic endemism appear to be patchier in past time horizons with a small region in the Canadian Maritimes arising as a location of high endemism (Figure 2C and 2D). Extending the idea of hotspots to areas gained and lost reveals strong decreases of the extent of ranges in the lower Rocky Mountains and Central Europe (Figure S2).

**Figure 2.**
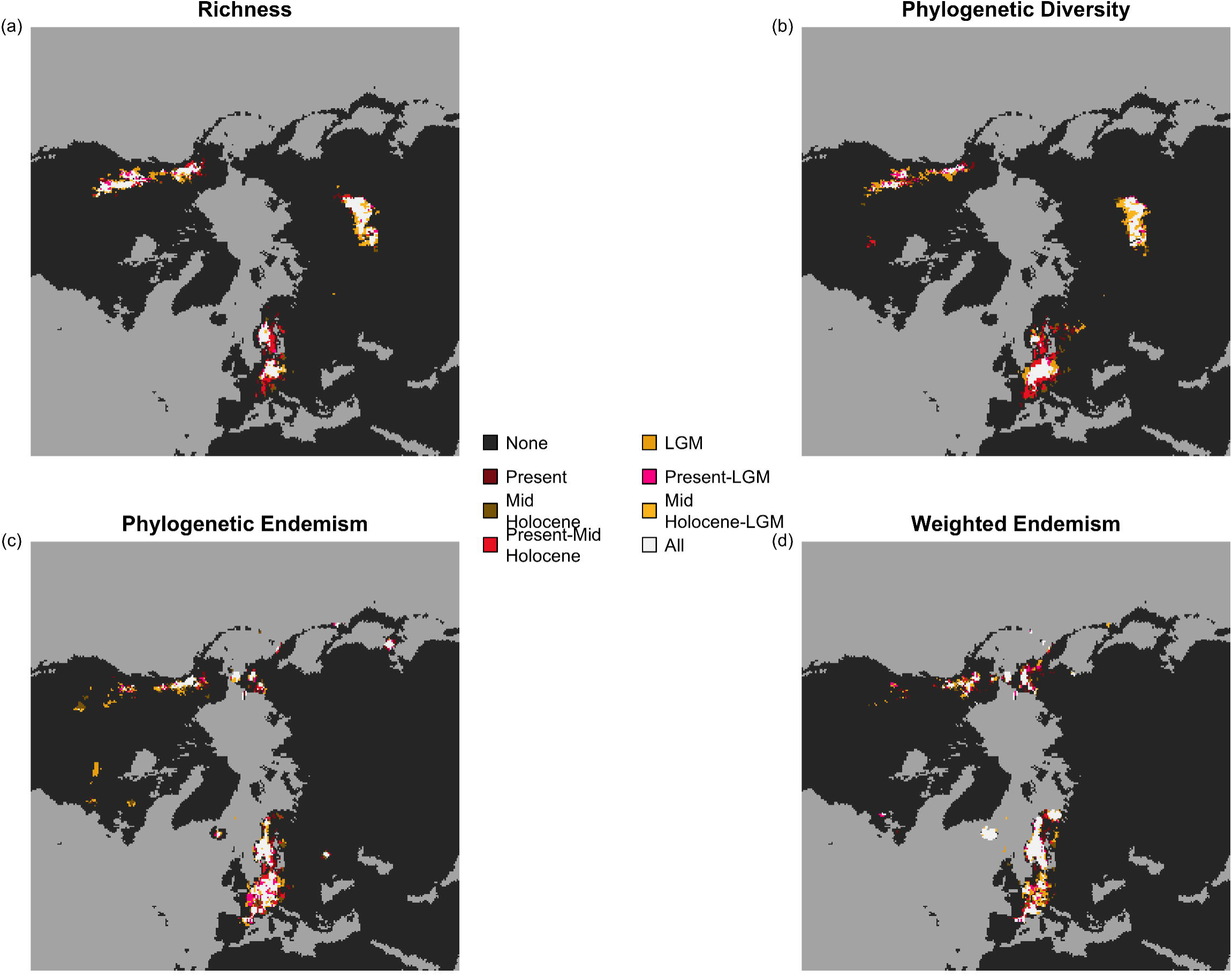
Hotspots of arctic alpha diversity. In each panel red colors correspond hotspots found in the present, while yellow colors were past-only hotspots. Dark gray is the background land area, while lighter gray is ocean. (a) Richness of the past and present. (b) Faith’s Phylogenetic Diversity. (c) Phylogenetic endemism. (d) Weighted endemism. Maps are shown in EPSG:3996 projection system.

### 2. Velocity and direction of range shifts

Beyond, range size and geographic shifts, we also investigated the velocity of shifts defined as the geodesic distance traversed by a species centroid between two time periods divided by the time spanned by the two periods (in km yr^−1^). When aggregating all plants, arctic species collectively moved either west or east, with a limited amount of movement north or west (Figure 3). Looking at just movement along the North-South axis, the Present-Mid-Holocene (median = 0.31 km yr^−1^+/- 0.02 km yr^-1^) and Mid-Holocene-LGM (median = 0.28 km yr^−1^+/-0.02 km yr^-1^) velocities indicated northward shifts across all plants while Present-LGM showed a southward shift (median = 0.32 km yr^−1^+/-0.02 km yr^-1^). The movements when comparing across times showed no significant difference in directionality between any of the three scenarios (Watson’s two sample test: U = 0.0342, p > 0.10). However, all centroid movements within each time horizon were significantly different from a uniformly circular distribution, indicating directionality in the data presumably along an East-West axis as opposed to a North-South one (Rao’s spacing test: U = 170.92, p < 0.001). Considering only along a scaled and absolute north-south velocity axis, we found that there were significant differences in the velocities by direction for the LGM-Mid-Holocene and LGM-Present (ANOVA for the LGM-Mid-Holocene movements: F = 2.612, p < 0.001, df = 1510; LGM-Present F = 4.06, p < 0.001, df = 1510), but not for the Mid-Holocene to the present (F = 1.278, p = 0.258, df = 1510) (Figure S2). Examining the differences closer using a Tukey’s Honestly Significance Difference test, we found that for the LGM-Mid-Holocene only movements in the north versus west (CI = (-0.62, - 0.11), p = 0.0003) and north versus east (CI = (0.026, 0.529), p = 0.0187) were significantly different from the others while southeast was only marginally significantly different to the Present-LGM (CI = (-0.0008, 0.50), p = 0.051) (Figure S3). Plant velocity directionality falls along an east-west axis as opposed to a north-south one as observed in lower latitudes. Mapping the shifts geographically, we found no significant difference in the velocities of species with centroids that moved across continents and those that stayed within in any comparison of the time horizons (example, Present-LGM F = 0.0197, p = 0.657). We lastly examined if there was a phylogenetic signal in the tendency of closely related species to shift at similar velocities than random expectations, we found significant phylogenetic signal for the Present-LGM (Pagel’s λ = 0.19, p = 0.026) and Present-Mid-Holocene (Pagel’s λ = 0.23, p = 0.001), indicating that species more closely related tend to move with the same velocities (Figure 4). However, there was no phylogenetic signal for the Mid-Holocene-LGM (Pagel’s λ = 0.04, p = 0.137).

**Figure 3.**
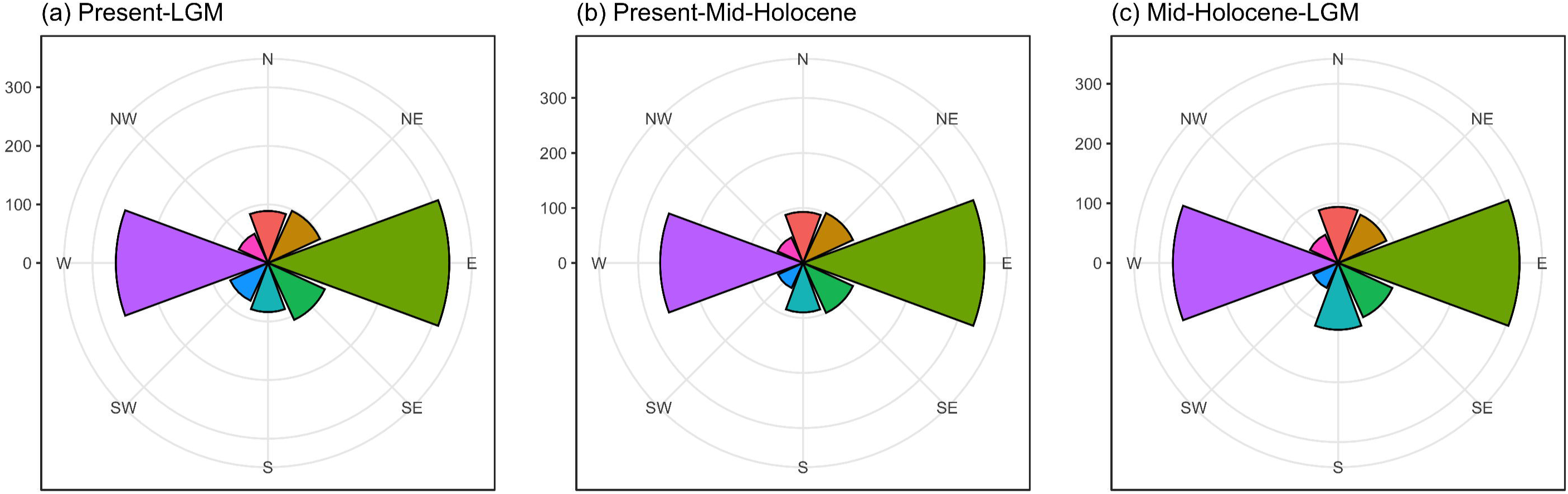
Direction of species centroid movement. (a) Present centroids versus centroids at the Last Glacial Maximuim. (b) Present centroids versus centroids at the Mid-Holocene. (c) Mid-Holocene centroids versus centroids at the Last Glacial Maximum. In all three comparisons the dominant movement was along the East-West axis, however only panel (b) the Present-Mid-Holocene was significantly different to random directions based on a Rao’s spacing test and none of these scenarios had movements significantly different from each other.

**Figure 4.**
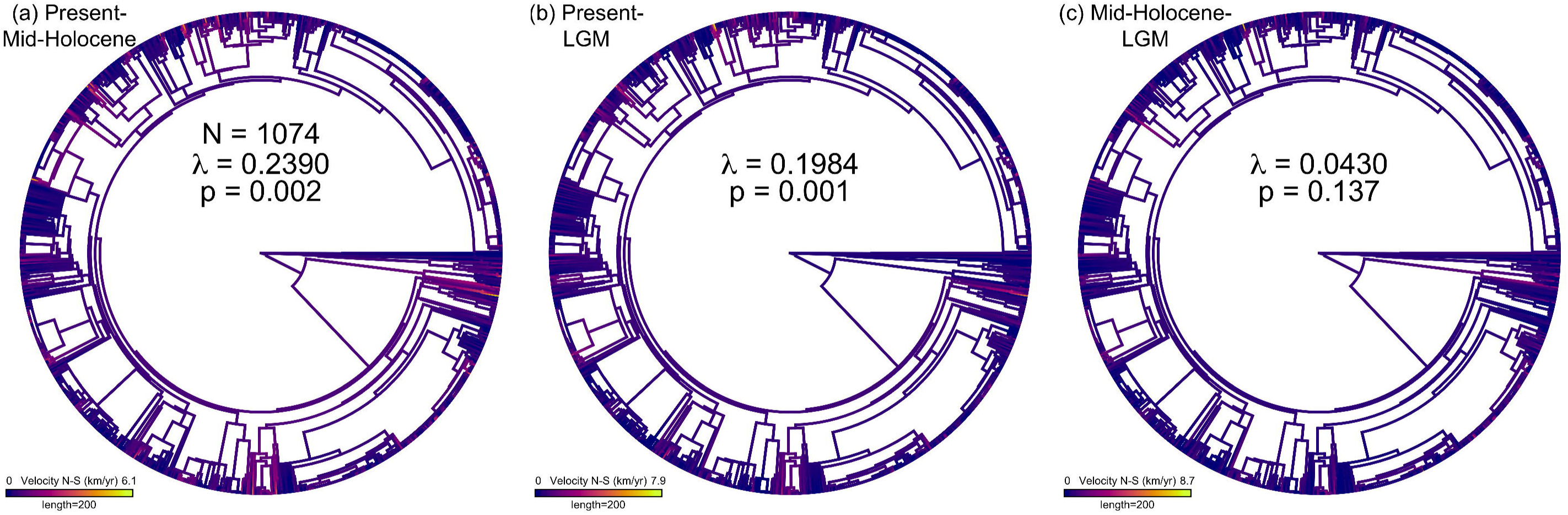
Phylogenetic patterns of species velocities along a north-south axis. Pagel’s lambda was significant for (a) present-midholocene and (b) Present-LGM but not for (c) Mid-Holocene LGM. Brighter colors indicate faster movement.

When considering elevational changes of individual plant species from the species distribution models, while we found a tendency of plants moving toward higher elevations at a velocity of 103 m yr^-1^ +/- 57.6 m yr^-1^ (Present-Mid-Holocene rate, 117 plant species) only 116 plant species had elevational shifts that were significantly different from their present-day ranges moving either upslope or downhill for the Present-LGM scenario at 228m yr^-1^, while the Present-Mid-Holocene had 119 at 184 m yr^-1^ (Figure S5). Of these, only 33 species (including Tofieldia coccinea, Cerastium articum, and Saussurea alpina) moved significantly either upslope or downslope in all three scenarios with most having an elevation significantly larger than expected under a null distribution. On average, most species moved upslope across the arctic (Figure S5D, E, F).

### 3. Validation of our predictions

We validated our predictions in three ways: (1) rerunning our models but excluding areas covered by the glaciers, (2) projecting past modeled predictions back to the present and, (3) by training our modeled estimates using Random Forest as a function of mammal richness whose diversity is well-known, environmental factors, and assuming a scenario of unbiased plant sampling.

We created reverse predictions where we used the outputs of the past modeled estimate as the input occurrences for modeling the present distribution assuming one were to do time travel back in time and project species distributions into the future being present day. This secondary run created fewer modeled outputs of the same species than in the original run, leading to widespread decreases in all four metrics when comparing between the original and reprojections (Figure S7). We ran the same comparisons as in section 1 of the results but compared against the original present and found that based on the modified t-test the two modeled presents were statistically different from the original present for all diversity metrics (Present vs Mid-Holocene-Present: F = 5.29, p < 0.001; Present vs. LGM-Present: F = 5.42, p < 0.001). In addition, when considering hotspots there are greater incongruities between the hotspots, notably in the phylogenetic and weighted endemism hotspots where far more patchy areas of high endemism are found in eastern Canada (Figure S8). These results do not necessarily imply that the validation failed but rather it may be an artifact of the lower total species richness of the reprojections. While the decreases observed in the alpha diversity metrics generally disagree with the original output, the hotspots and values nearby the hotspots remain roughly the same, indicating the validity of this method for detecting hotspots.

We then modeled species distributions during the LGM but excluding areas covered by the glaciers at the height of the LGM. This was done as above and like above, there were fewer secondary modeled output vector polygons than in the original, most likely due to memory segmentation errors while running the MaxEnt model (Figure S9). This is reflected when taking the percent difference between the two scenarios. When considering the hotspots of diversity metrics for species distributions during the LGM, both the glaciated and unglaciated scenarios appear to have good alignment for almost all metrics (Figure S10).

Lastly, we validated our results by training a random forest on the modeled estimates as a function of climatic variables with the addition of tetrapod richness, a group whose geographic distribution is well-known, and under a scenario of high sampling of occurrences. We found strong alignment between the predictions based on the original modeled estimates versus Random Forest, given by the large overlap between the two in their hotspots (Figure S11).

## Discussion

We present an evaluation of historical changes of plant diversity patterns in the Arctic since the Last Glacial Maximum and mid-Holocene using species distribution modeling as a function of present-day species occurrences. Our findings indicate (1) divergence in spatial patterns of alpha diversity with increases typically between the Present to LGM and decreases from the Mid-Holocene to the Present; (2) consistence in the direction and velocity of species movement but are not phylogenetically patterned; and (3) elevational shifts in arctic plant species are limited to a small number of species. We elaborate on these findings below.

Although each prediction at each time point was significantly different from the other time periods, the overall change in richness, phylogenetic diversity, phylogenetic endemism, and weighted endemism do not appear to show a clear and coherent signal whether increasing or decreasing globally save for the Mid-Holocene to Present transition. We also find in our models higher species richness at the southern edge of the arctic, along the boreal forest-tundra ecotone. We identify the Rocky Mountains, Inner Mongolia and Central Europe as hotspots of richness and phylogenetic diversity, and Iceland and Far East Siberia as hotspots of endemism for arctic plants. The hotspots in the Rocky Mountains and Central Europe would be expected as the maximal extent of the LGM glaciers stopped close to it [7. Molnar et al. 2023; 8. Beatty et al. 2010]. It is possible that arctic plants likely occupied this region during the early phases of retreat and then spread outward into the Alps and Scandinavia. The distributions of plants in the arctic is determined by when deglaciation occurred as evidenced by previous studies that found that landscape age strongly influenced species richness and genetic diversity, which is consistent with what we present here [1. Stewart et al. 2016; I2-3]. Other studies [e.g., 2. Zemlinsky et al. 2024] found that past climatic conditions from the Mid-Holocene and LGM strongly influenced the present richness of arctic plants. Although not explicitly stated as a biodiversity hotspot, the literature suggests that as a refugia for plants, Beringia should have relatively high plant diversity, which is consistent with what we found for endemism. For species range sizes, we show that most species have lost their ranges in the Rocky Mountains and Central Europe, likely due to the reduction in hotspot extent in these regions, while most gains happened in Far East Siberia, aligning with the hotspot of endemism present there. Such findings could indicate that even over longer periods, the changes in arctic alpha diversity confirm previous findings that no trends in richness or community composition despite observed warming across the bioclimatic delineation of the arctic using vegetation surveys over 40 years [3. Garcia-Criado et al. 2025].

The directionality of the movement of plants remained consistent across all three time horizons, meaning that the movement in any scenario was ultimately similar across time horizons. The lack of a geographic signal between time horizons is reflected when plotting this data geographically as the mean direction by cell appears to be nearly random. In addition, the movement along this axis holds across phylogenetic groups indicating that all plants move consistently like this. When considering velocities of species centroid (center of distribution) movement only along a north-south axis the direction of movement appears to separate along a northwest-southeast axis as opposed to a strictly north-south one. Our study diverges from previous published literature on the direction of range shifts. For instance, while some studies document northward shifts in plant ranges at lower latitudes [e.g., 4. Van der Putten 2012], we find instead that most arctic plant species shifted their ranges either east or west. For velocities, we found a strong phylogenetic signal in Pagel’s λ for every scenario indicating that species velocities in response to climate change can be partly attributed to evolutionary history. We note several clades in Caryophyllaceae that appear to have elevated velocities in the North-South direction (Figure 4). Taxa in this family are far more likely to shift ranges either north or south at a much faster rate than expected under Brownian motion, possibly due to their shared evolutionary history which for those adapted for cold temperatures result in unspecialized seed dispersal [9. Tovar et al. 2020]. Similarly, we find a restricted number of species shifted their mean elevational range upwards, while at least one shifted significantly lower in elevation. The lack of a phylogenetic signal with respect to elevational shifts may be in part because most of the arctic is relatively flat, topographically. Higher elevational sites that could be occupied require far greater dispersal to the lower Rockies from the Canadian Archipelago or to the Alps from Siberia or even the Ural Mountains. While all arctic plants appear to consistently remain within the arctic based on the east-west centroid shifts, only a few plant clades are shifting their ranges quickly north-south and fewer still are shifting their ranges higher in elevation.

Our findings here support a hypothesis on species distributions of arctic floral biodiversity in that despite a warming climate there is limited evidence of strong changes in alpha diversity, namely species richness [3]. However, there is a great contrast between the patterns that occur in the arctic and those at lower latitudes, namely the stark departure from previously established patterns of northward and upward range shifts at lower latitudes [4; I-31]. Plants in the arctic are already at the farthest terrestrial north possible to occupy, so it is possible that they continually shift ranges east-west as aided by long-distance dispersal [I-2]. This may explain the lack of a clear increase or decrease in many of our alpha diversity metrics (Figure S1) and the persistence of large hotspots throughout our modeled time points (Figure 2). Alpha diversity in the arctic is different between time horizons in this study, as well as to the differences in alpha diversity, but we could not conclude whether it is increasing or decreasing, only that it is different.

While there are limitations in the interpretation of the results of this study, namely in the ability to ground-truth the data with fossil data, the validation methods used here support our discussion. Some known drivers of arctic plant distributions are not included in the SDMs for this study, namely those related to snow [5. Niittynen et al. 2018]. The lack of inclusion of variables related to snow such as snow cover duration and melt timing are caused primarily from a lack of readily available data on the global scale and at the time horizons considered in this study. In addition, because we did not examine compositional turnover, we may have missed changes in species composition across the landscape over time. This study is limited in that there is a lack of readily available fossil resources to ground truth these modeled estimates. Fossil data are generally sparse for identifying individual plant species presence [6. Aarnes et al. 2012]. In addition, plant sampling in the arctic is prevalent with sampling biases and gaps given how remote and sparsely populated the arctic is, leading to a greater density of observations coming from roadsides and urban centers [I-3, I-27]. Lastly, one of the major hinderances to accurately describing species distributions at any time point is due to the lack of information readily available on species interactions. Here, we assumed that only abiotic factors drive distributions. However, our results from the random forest analysis of richness support our present findings in that it is an assessment independent modeled estimates based on MaxEnt and assumes both equal and unequal sampling across the arctic in the two scenarios we analyzed.

We present here a picture of plant range changes and elevational changes since the LGM in the arctic. The arctic is a very dynamic system that although may have a lower number of species given its large expanse, it still presents a fascinating opportunity to understand global swings in temperature both seasonally and over long time periods. With our results, especially for those identifying biodiversity hotspots, it is possible to potentially target certain areas for conservation of areas with both high plant endemism and diversity in the past and present. In addition, our finding of a phylogenetic signal for species velocities for species in the present may assist conservationists and future studies on arctic flora by providing groups of taxa known to collectively respond more strongly to climate change. Far more research is needed to fully capture the biological mechanisms of plant establishment and persistence in this extreme and rapidly changing environment.

## Data Availability Statement

Species occurrence data is available from GBIF (https://doi.org/10.15468/dl.t7mpsk). Climate data can be downloaded from worldclim.org. Phylogenetic data taken from https://github.com/FePhyFoFum/big_seed_plant_trees/releases. Input data and species distribution models are available from Data Dryad (https://doi.org/10.5061/dryad.zkh1893p2.)

Reviewer Link: http://datadryad.org/share/rFzLpLg68HxFx8Od_FBmHoRI3Kq4N3Wc9K4Ffo5C9vo

---

**Figure S1.**
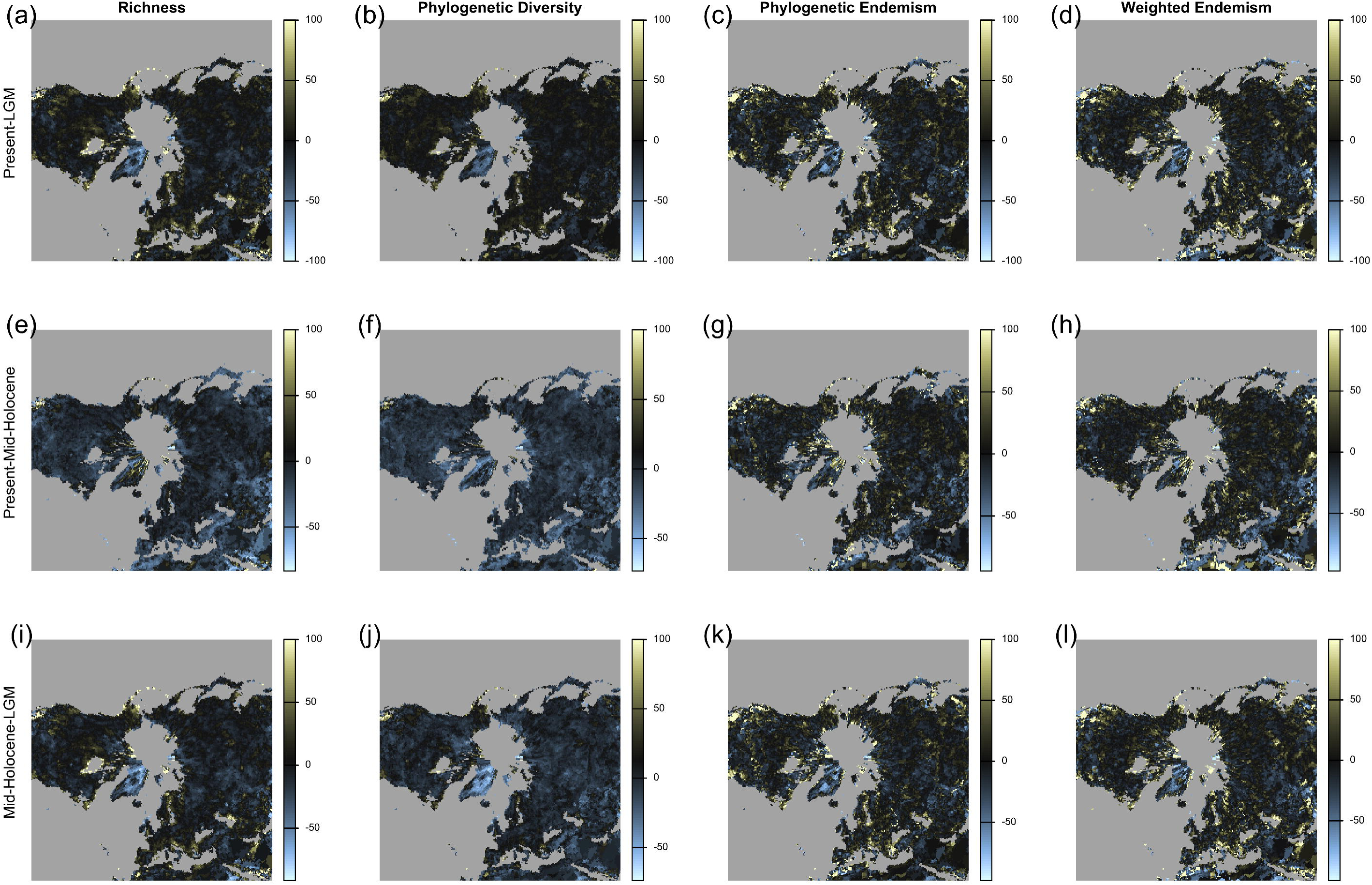
Changes in plant alpha diversity based on species richness, phylogenetic diversity, phylogenetic endemism, and weighted endemism. a-d, alpha diversity metric change between the Present and LGM for (a) species richness, (b) Faith’s Phylogenetic Diversity, (c) phylogenetic endemism, and (d) weighted endemism; e-h, Present and Mid-Holocene (e) species richness, (f) PD, (g) phylogenetic endemism and (h) weighted endemism, and; i-l Mid-Holocene and LGM showing (i) richness, (j) PD, (k) phylogenetic endemism, and (l) weighted endemism.

**Figure S2.**
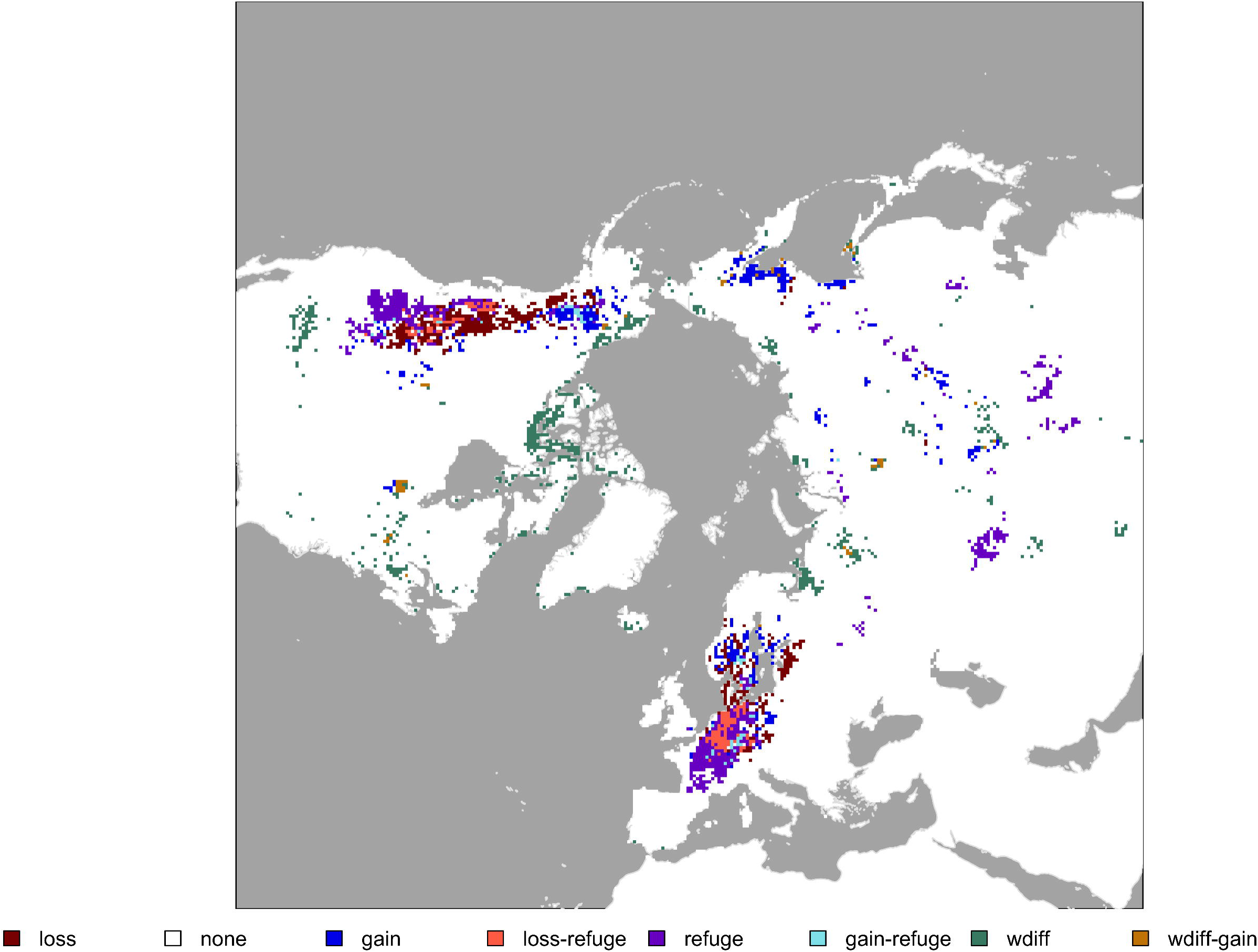
Hotspots of range size increases and decreases. Range expansions and contractions were determined through geometric operations by clipping between time horizons for each individual species. Areas of refuge were determined through intersecting range vector polygons across all three time horizons. The resulting vectors were then rasterized. The areas gained raster was subtracted by the areas lost raster and then divided by the refuge raster which we called weighted difference. All rasters were then thresholded at the 97.5^th^ percentile to determine hotspots of areas lost and areas gained. Map is projected in EPSG:3996 and grid cells are 50km by 50km.

**Figure S3.**
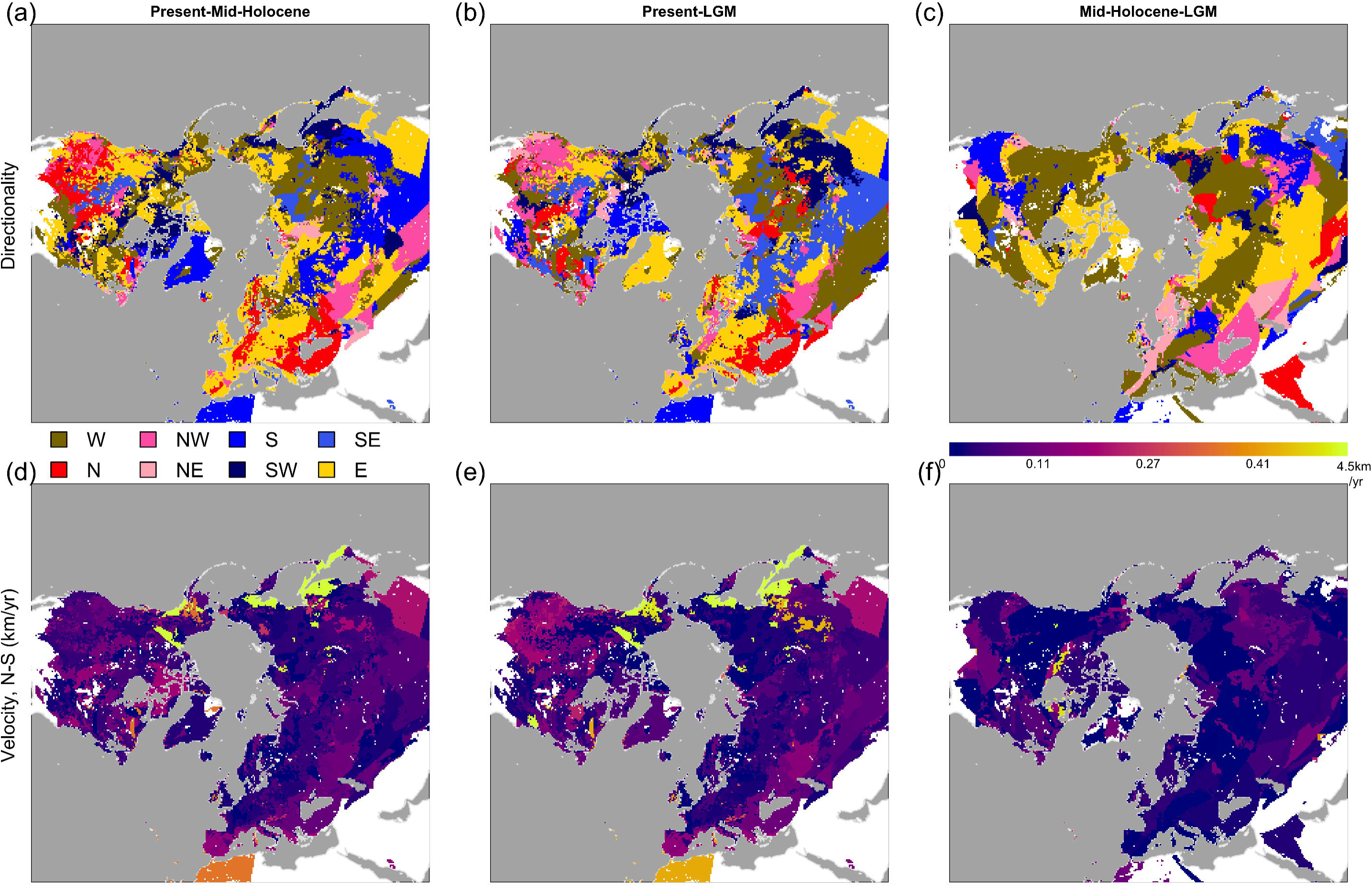
Directionality and velocity of species range shifts. (a)-(c), Aggregated direction of species centroid movements. (a) Present versus Mid-Holocene. (b) Present versus Last Glacial Maximum. (c) Mid-Holocene versus Last Glacial Maximum. Grid cells are sized at 50 km by 50 km, legend is in the middle left. (d)-(f), velocity of species centroid movement along a North-South axis. (d) Velocities of the present using the Mid-Holocene as reference. (e) Velocities of the present using the Last Glacial Maximum as reference. (f) Velocities of the Mid-Holocene using the Last Glacial Maximum as reference. Legend is in the middle right. Maps projected in EPSG:3996.

**Figure S4.**
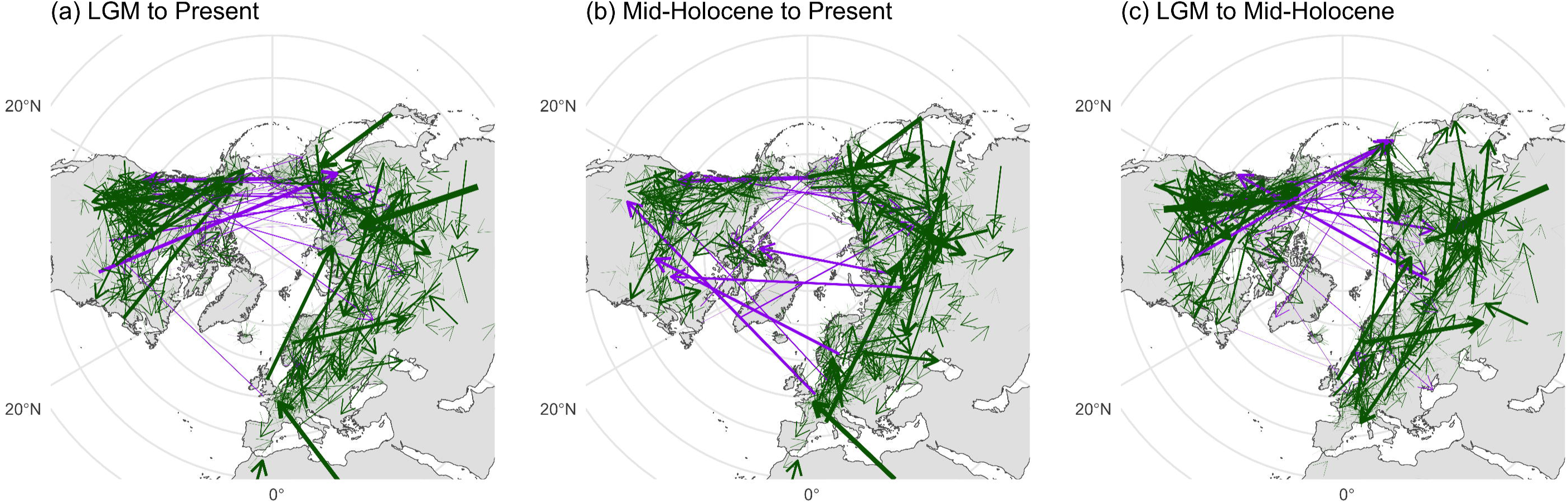
Centroid movements across the arctic. Green arrows correspond to centroid movement within biogeographic realms, or in this case, within continents while purple arrows indicate movements across continents. Not pictured are centroids that either started or ended over water. One arrow is indiciative of one species alone. (a) Movement of centroids from the Last Glacial Maximum to the Present. (b) Movement of centroids from the Mid-Holocene to the Present. (c) Movement of centroids from the Last Glacial Maximum to the Mid-Holocene. There was no significant difference between the velocities of centroids that moved across continents and those that did not in any time horizon. Arrow size is proportional to centroid velocity.

**Figure S5.**
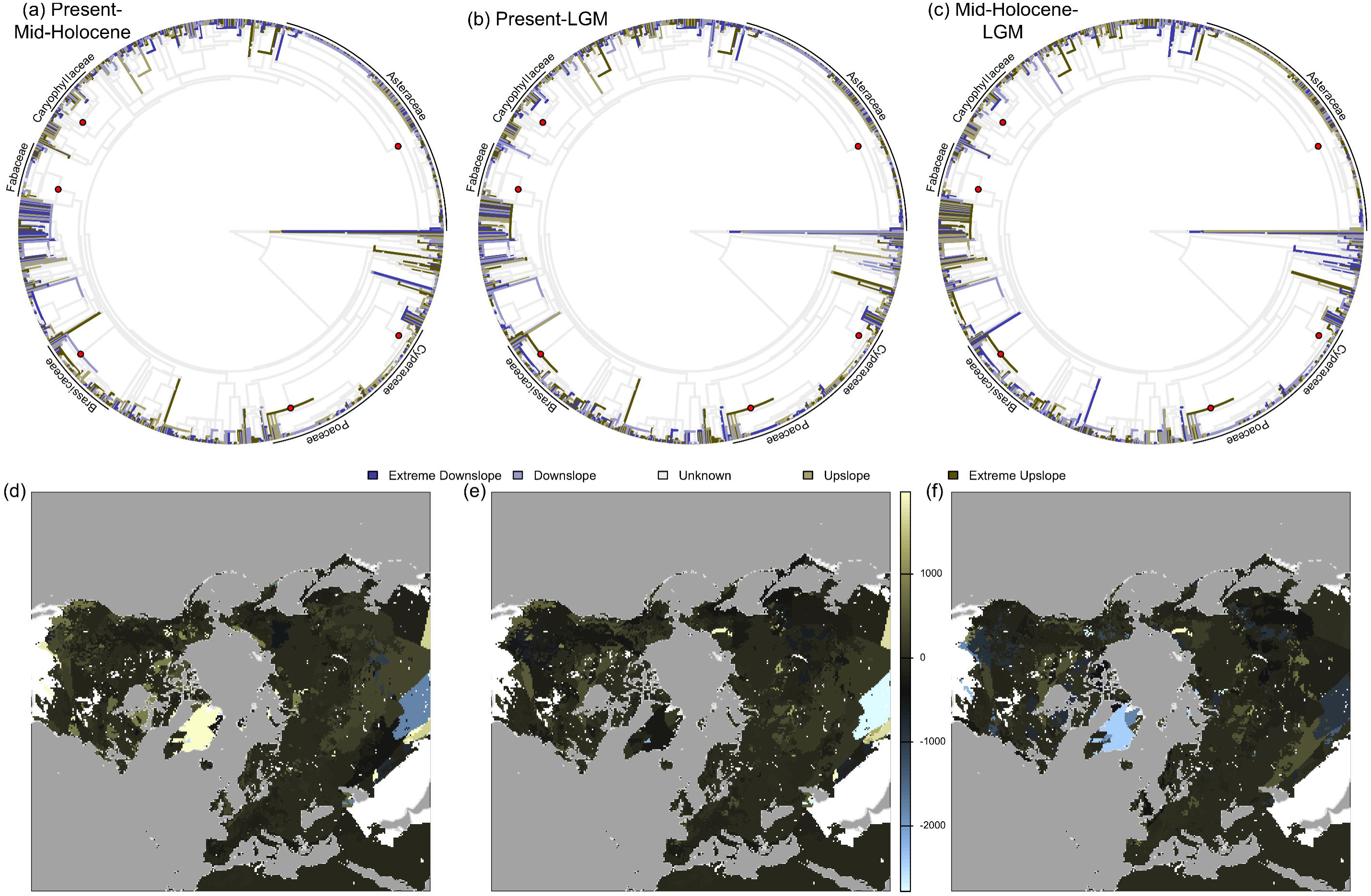
Phylogenetic and geographic elevational shifts. Panels A-C, elevational shifts binned into four categories. Extreme downslope was -1000m to -400m, downslope was -400m to 0, upslope 0 to 400m, and extreme uplsope was 400m to 1000m. Represented in the phylogeny are 1511 species; there was no phylogenetic signal following Pagel’s lambda in any of the scenarios. Panels D-F, geographic aggregation of species elevational shifts. Most species moved slightly upslope, but not significantly so. Maps projected in EPSG:3996.

**Figure S6.**
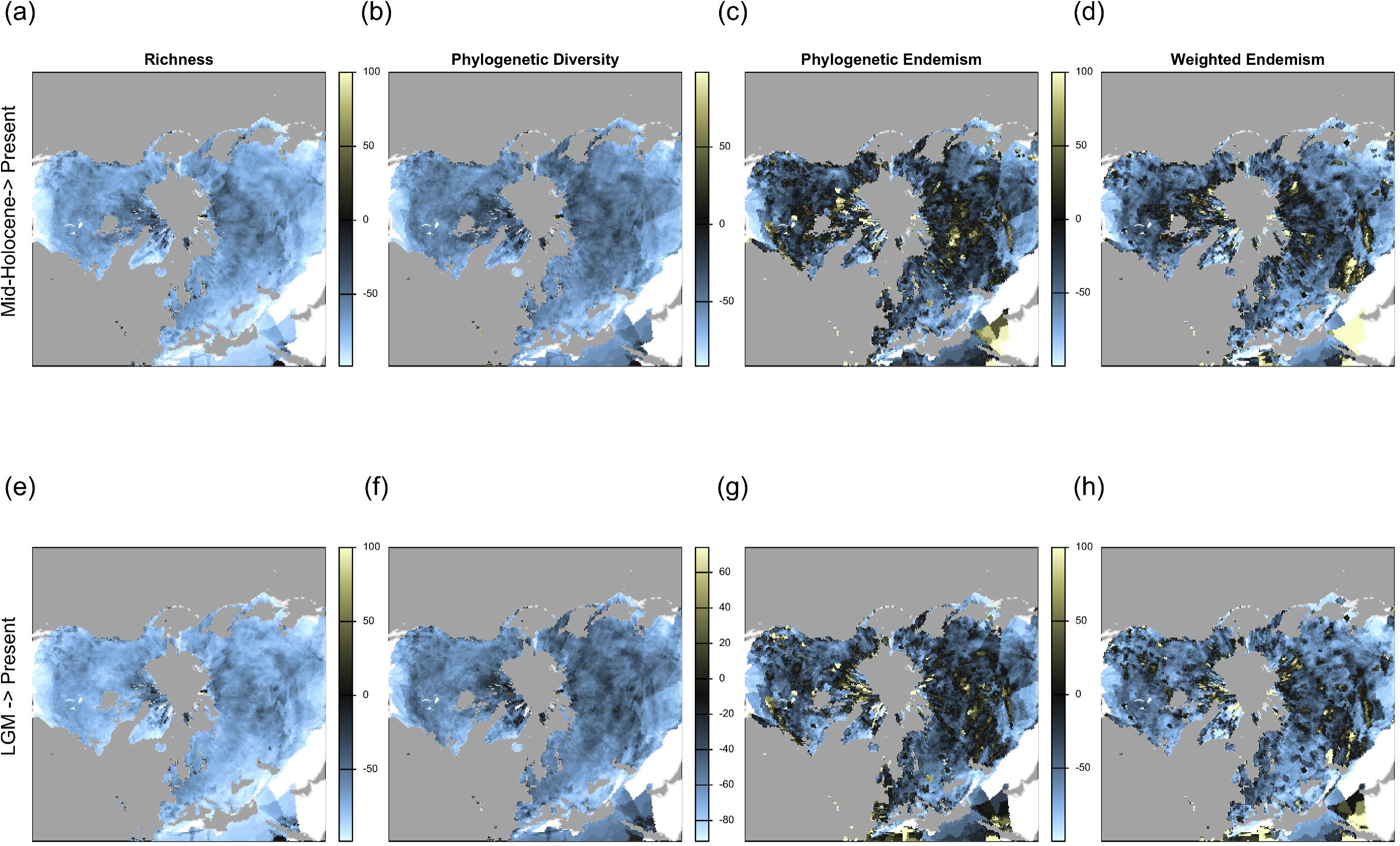
Differences between original Present SDM and reprojection of past scenarios to the present. (a) – (d) Richness, Faith’s Phylogenetic Diversity, Phylogenetic Endemism, and Weighted Endemism when projecting past SDM outputs to the present. The lower row shows projections of the Last Glacial Maximum (LGM) back to the present for (e) Richness, (f) Faith’s Phylogenetic Diversity, (g) Phylogenetic Endemism, and (h) Weighted Endemism. For both cases, Richness and phylogenetic diversity showed increases outside of hotspot areas. Both endemism metrics reported increases, likely due to decreased range sizes and reduced richness.

**Figure S7.**
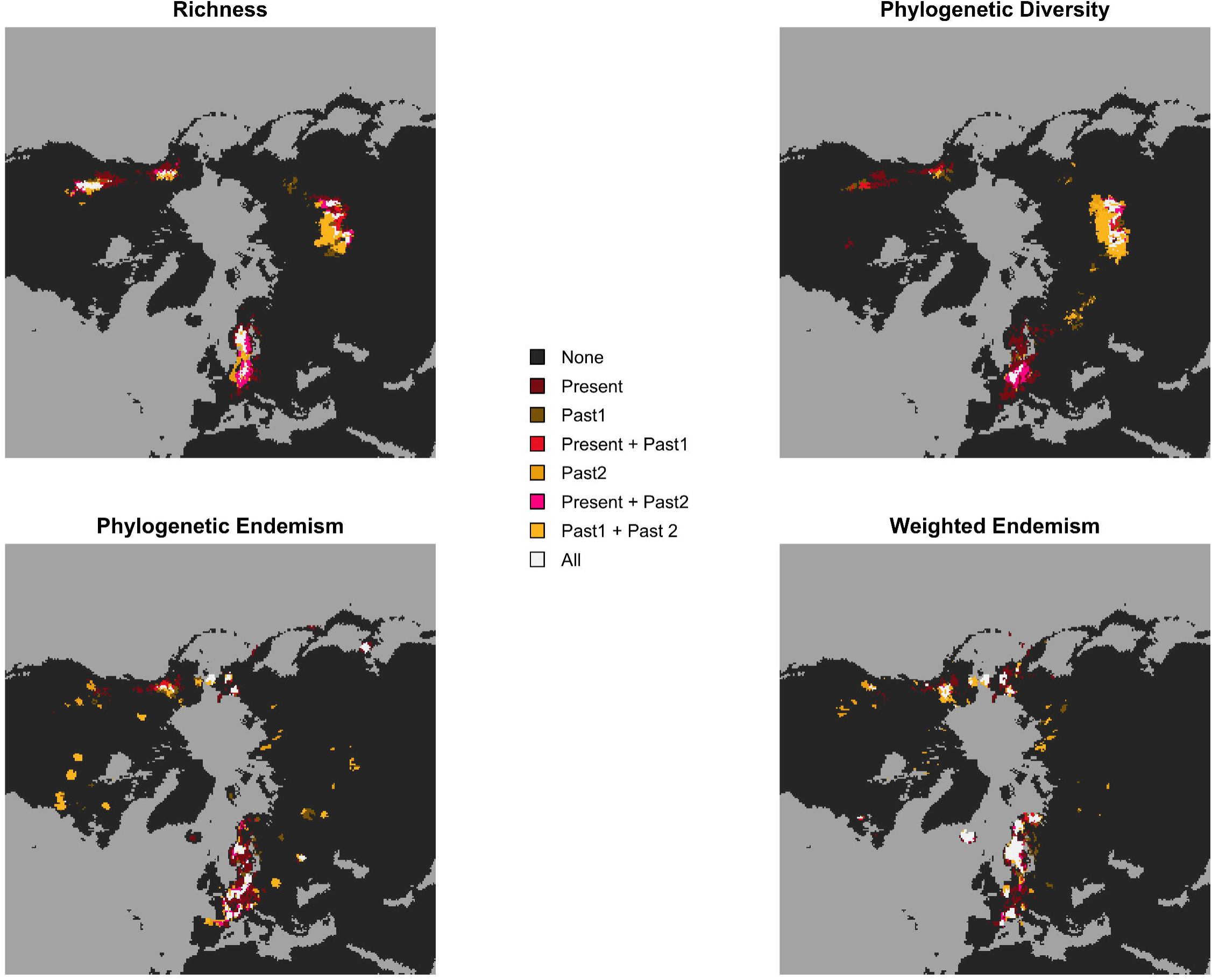
Hotspots of past reprojections to the present. Red colors indicate reprojection agreement with the present. White shows agreement across both reprojections and the present, while yellow shows only reprojection hotspots. There is major disagreement in Mongolia for richness and phylogenetic diversity, while agreement in central Europe for all scenarios.

**Figure S8.**
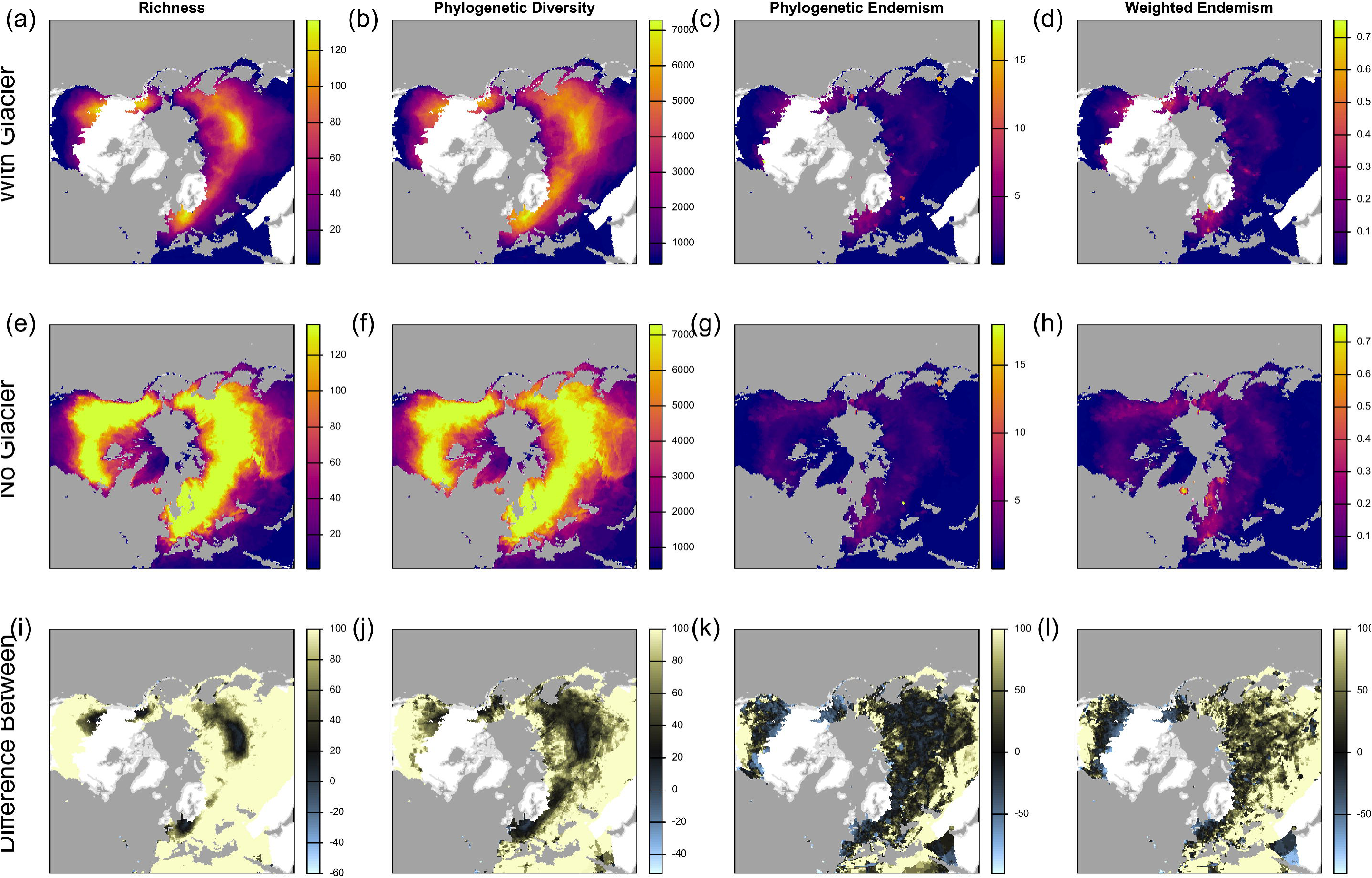
Comparison of alpha diversity metrics when exlcuding land covered by the glacier. Metrics for the LGM-Glacier are consistently lower than in the original LGM. There is however agreement, mostly in areas previously identified as hotspots. Top row presents predictions for the LGM time horizon without allowing for presences in areas putatively covered by the glaciers for (a) Richness, (b) Faith’s Phylogenetic Diversity, (c) Phylogenetic Endemism, and (d) weighted endemism. Middle row is the same as presented in figure 1, showing (e) species richness, (f) Faith’s PD, (g) phylogenetic endemism, and (h) weighted endemism for the LGM allowing for presence in any terrestrial cell. Bototm row shows the difference between the top row and middle rows for each of (i) species richness, (j) PD, (k) phylogenetic endemism, and (l) weighted endemism.

**Figure S9.**
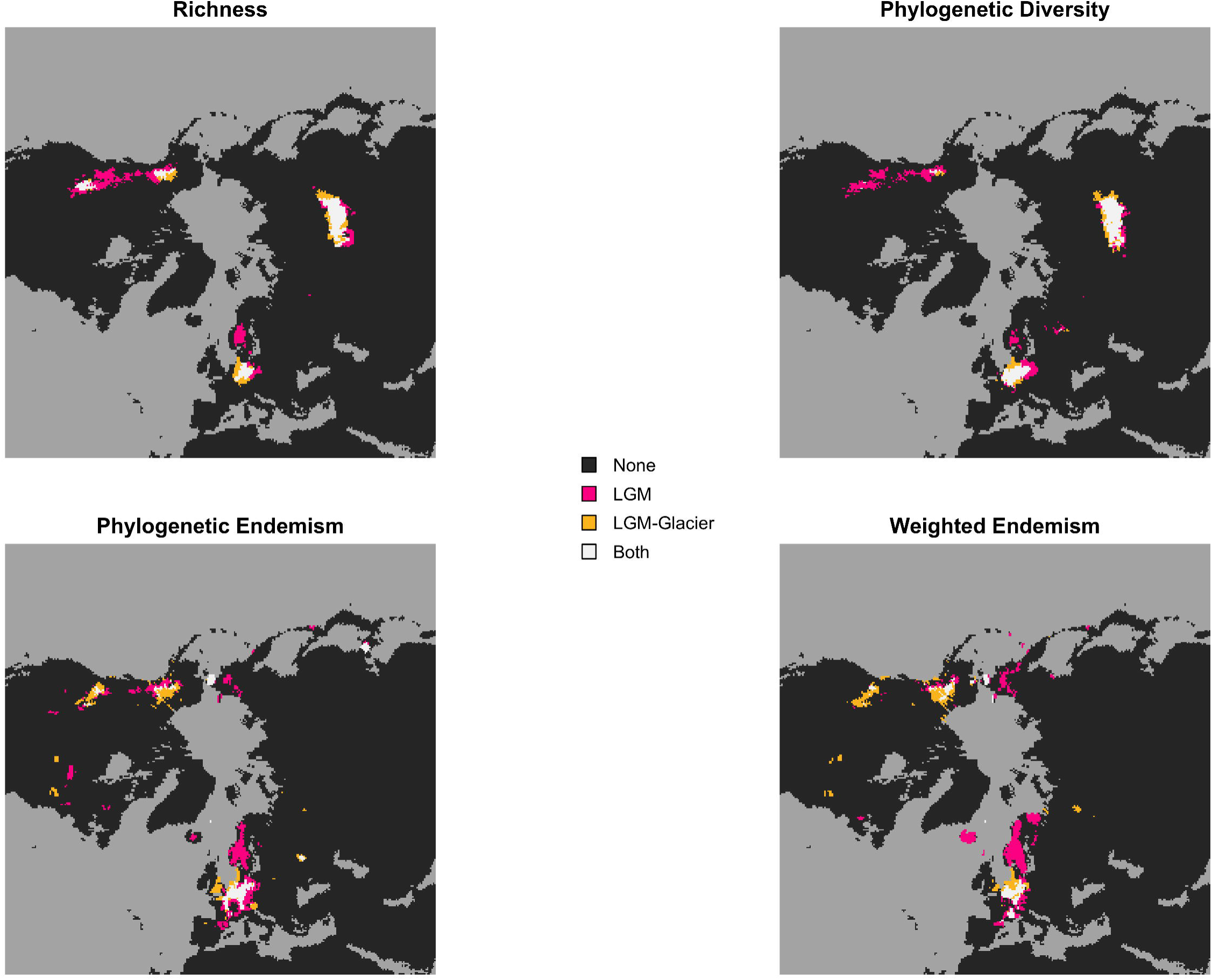
Comparison of hotspots between the original LGM prediction and the LGM-Glacier prediction. Mongolia and Central europe show good alignment between the two scenarios, however the divergence between the two becomes more apparent when looking at the Rocky Mountains and Scandanavia.

**Figure S10.**
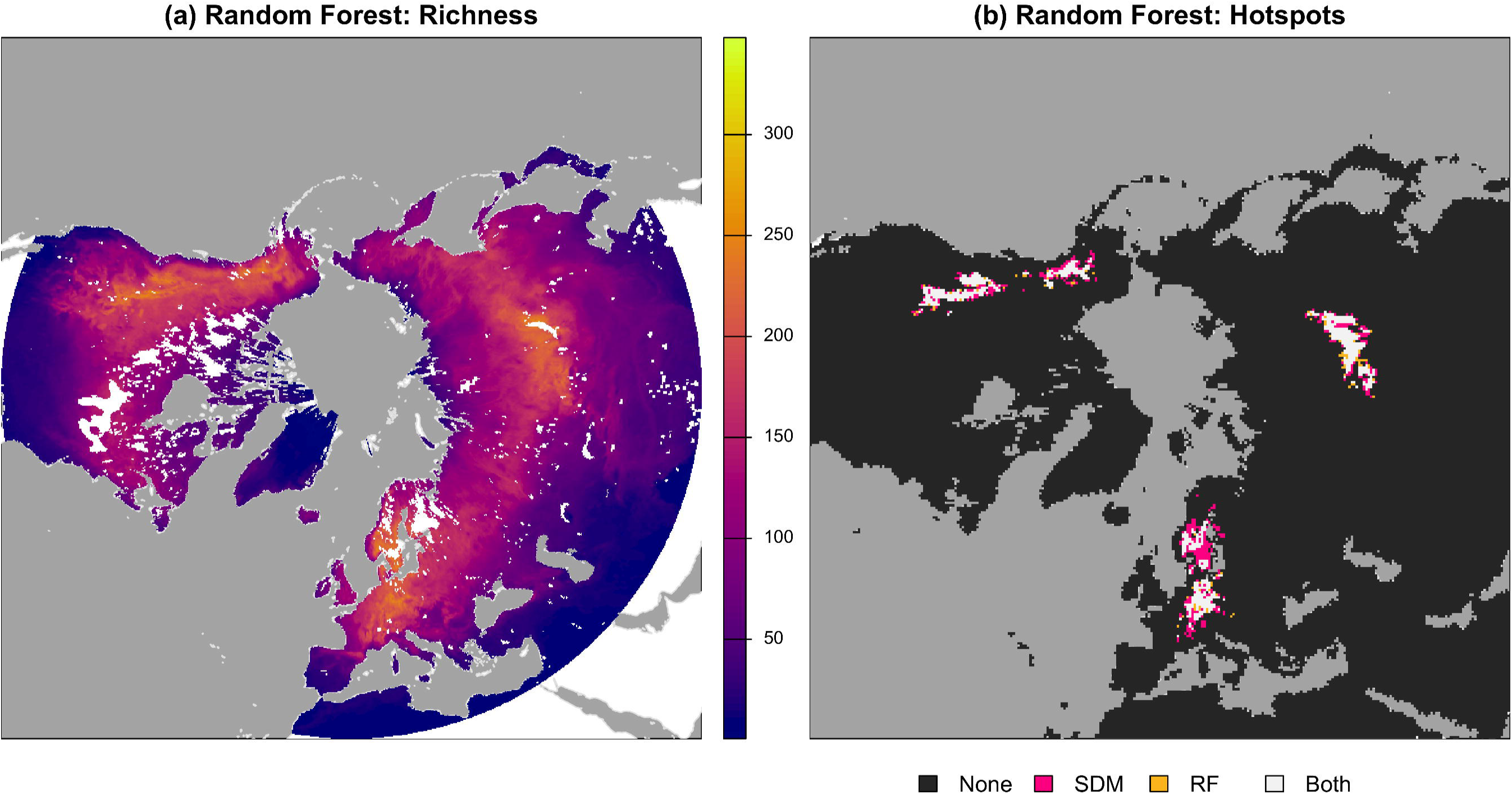
Random Forest Richness predictions and hotspots agreement with MaxEnt. Shown here in (a) is the random forest output for unbiased sampling with the inclusion of tetrapod richness. (b) hotspots of richness between the two models align well, as indicated by the large amounts of white. MaxEnt appears to predict relatively larger hotspots than random forest.

**Figure S11.**
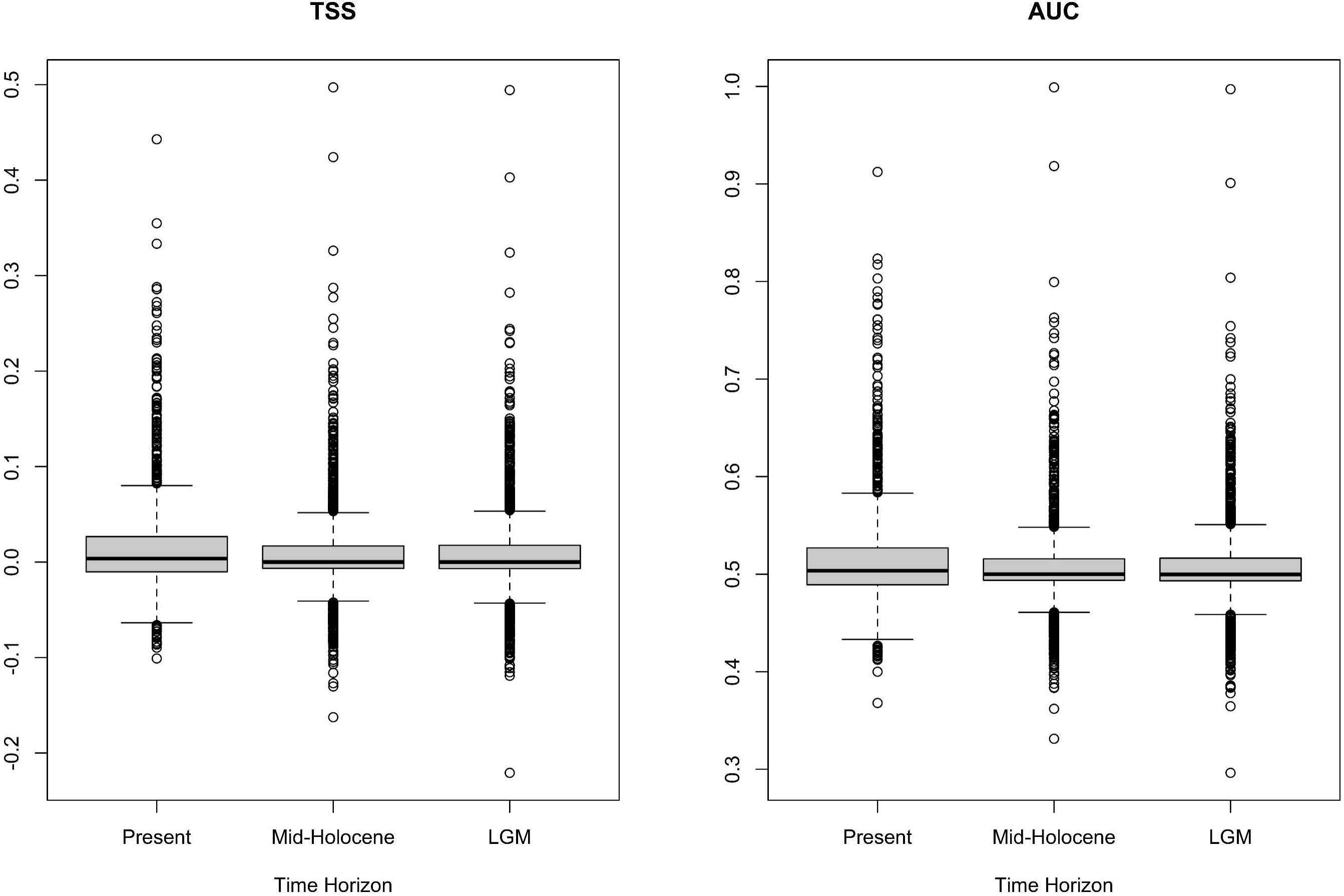
Model validation results. (a) True Skill Statistic and (b) Area Under the Curve for the three modled time horizons.

